# Dispersal heterogeneity, rather than dispersal evolution, drives biodiversity patterns

**DOI:** 10.1101/2025.11.27.690991

**Authors:** Siebe van Wunnik, Felipe Kauai, Dries Bonte

## Abstract

Island biodiversity emerges from a fundamental tension between colonization and divergence. Dispersal enables lineages to reach new habitats and exploit ecological opportunity, but simultaneously increases gene flow that can inhibit the genetic differentiation required for speciation. Although dispersal is widely recognized as a key driver of biodiversity patterns, most theoretical studies treat it as a fixed species attribute, making it difficult to distinguish the effects of dispersal evolution, standing dispersal variation, and the spatial genetic structure generated by dispersal inheritance.

Here, we develop a spatially explicit individual-based model in which dispersal is a heritable quantitative trait and compare three dispersal scenarios: fixed dispersal, heritable evolving dispersal, and dispersal reshuffling that maintains trait variation while removing local inheritance. Across structured (archipelago-like) and unstructured landscapes, we evaluate how these different components of dispersal shape colonization, diversification, and long-term community dynamics.

We find that dispersal heterogeneity is a stronger determinant of biodiversity than dispersal evolution itself. In structured landscapes, both evolving and reshuffled dispersal substantially increase species richness, spatial turnover, and island colonization relative to fixed dispersal, whereas in unstructured landscapes dispersal variability reduces richness. Speciation and extinction rates remain closely balanced across scenarios, indicating that differences in diversity emerge primarily through changes in colonization and spatial organization rather than through altered diversification rates. When dispersal evolves, newly formed species follow two distinct trajectories, characterized by either increasing or decreasing dispersal capacity after speciation. Contrary to expectations from island-syndrome theory, the relative frequency of these trajectories is only weakly associated with landscape connectivity.

Our results show that individual variation in dispersal capacity fundamentally alters diversification dynamics by creating opportunities for both colonization-driven and reduced-gene-flow pathways to speciation. Incorporating dispersal heterogeneity into island biogeography therefore provides a more mechanistic framework for understanding how biodiversity emerges and persists in spatially structured landscapes.

## Introduction

Dispersal modulates gene flow and ecological opportunity within populations, acting both as a catalyst and a constraint in the formation of new species. High dispersal enables individuals to reach novel habitats, escape competition or predation, and exploit ecological opportunity, thereby supporting divergence and adaptive radiation (Cowie & Holland 2006; Losos & Ricklefs 2009; Lorion et al. 2013, Price & Wagner 2004). At the same time, dispersal facilitates gene flow, homogenizing genetic variation and counteracting the differentiation required for reproductive isolation (Bohonak 1999; Hendry et al. 2001; Givnish 2010). This duality generates a fundamental paradox: the process that enables colonisation can also inhibit divergence through increased connectivity. Understanding how dispersal shapes this balance remains a central challenge in evolutionary ecology (Hendry 2017; Pelletier et al. 2009; Schoener 2011).

Islands provide an exceptional context in which this tension is particularly pronounced. Their discrete boundaries, isolation, and limited area influence colonisation and extinction dynamics as well as ecological releases that differ sharply from continuous landscapes (Matthews & Triantis 2021). As a result, island populations frequently experience reduced competition and novel ecological conditions that promote predictable shifts in morphology, behavior, and life-history traits, collectively known as island syndromes (Adler & Levins 1994; Goltsman et al. 2005; Novosolov et al. 2013; Baeckens & Van Damme 2020; Jezierski et al. 2024). Among these traits, dispersal evolves repeatedly, with many insular species exhibiting reduced dispersal relative to mainland relatives (Carlquist 1966; Cody & Overton 1996; Fresnillo & Ehlers 2008; García-Verdugo et al. 2017; Jessop et al. 2018; Schenk 2013). These patterns underscore how spatial isolation shapes evolutionary trajectories, producing elevated levels of endemism and adaptive radiations (Givnish 2010; Yoder & Yang 2004), creating conditions in which the evolution of dispersal may determine whether lineages persist, diversify, or disappear.

Although theoretical frameworks for dispersal evolution are well established (Ronce 2007; Duputié & Massol 2013), further work is needed to understand how evolving dispersal feeds back into the processes of speciation and community assembly. This question is particularly relevant given evidence that dispersal is heritable, subject to natural selection, and responsive to ecological gradients and biotic interactions (Bonte et al. 2012; Saastamoinen et al. 2018; Duputié & Massol 2013; Fronhofer et al. 2024). Eco-evolutionary feedbacks involving dispersal influence priority effects (De Meester et al. 2016; Fukami 2015), gene flow (Alzate et al. 2019; Thompson & Fronhofer 2019), range dynamics (Van Petegem et al. 2018), and local adaptation (Hendry et al. 2001). Yet an important unresolved question is whether biodiversity patterns are driven primarily by dispersal evolution itself, by the resulting variation among individuals in dispersal capacity, or by the spatial genetic structure and subsequent speciation. Because these components are typically confounded, their relative contributions to colonisation, diversification, and extinction remain poorly understood. Although classical island biogeography theory predicts species richness through colonisation-extinction balance (MacArthur & Wilson 1967), it does not account for evolutionary feedbacks arising from evolving dispersal.

Understanding diversity patterns across the tree of life may thus require an explicit account of the generation substrate in which dispersal evolves. Genetic variance and its distribution over spatially concentrated populations provide a link between ecological processes and evolutionary outcomes. Standing genetic variation determines whether dispersal can respond adaptively (Barrett & Schluter 2008), while local genetic structure governs how dispersal feeds back on trait variation through gene flow, inbreeding, and adaptive differentiation (Lenormand 2002). Importantly, these two components need not have equivalent ecological consequences: dispersal variation can alter colonisation opportunities even when local inheritance is absent, whereas spatial genetic structure emerges from the persistence and transmission of that variation through space and time. Consequently, dispersal is an eco-evolutionary mechanism that redistributes genetic diversity, modulates local adaptation, and alters population persistence (Edelaar & Bolnick 2012; Garant et al. 2007). By altering connectivity and isolation, genetic variance and its spatial structure determine whether dispersal promotes evolutionary rescue and range expansion, or instead accelerates maladaptation, lineage divergence, and demographic collapse (Carlson et al. 2014).

The tension created by dispersal’s opposing effects on divergence is also central to a broader question in evolutionary ecology: which mechanisms maintain stable species richness over long timescales? Diversity may reflect either a static equilibrium, in which speciation and extinction are rare, or a dynamic equilibrium, in which turnover is high but balanced. Comparative studies suggest that multiple mechanisms can generate stable richness depending on ecological context and evolutionary pressures (Rabosky 2009; Morlon et al. 2010; Wiens 2011). Incorporating dispersal as an evolving trait in models of speciation may provide a more realistic and useful framework for properly distinguishing these alternatives.

To address these questions, here we develop a spatially explicit, individual-based model of speciation, in which dispersal evolves as a heritable trait. We compare this evolving scenario with scenarios where dispersal is held fixed (homogeneously distributed across individuals) or kept uniformly distributed across individuals over time (reshuffled), allowing us to isolate the role of genetic variance and spatial structure. This framework allows us to determine whether biodiversity patterns emerge from adaptive changes in dispersal, from variation in dispersal capacity itself, or from the spatial organisation generated by the inheritance of dispersal traits. By explicitly disentangling these mechanisms across structured and unstructured landscapes, we provide a mechanistic understanding of how dispersal shapes diversification, metacommunity dynamics, and the maintenance of biodiversity in spatially structured systems such as archipelagos.

More specifically, we ask:

(1) How do structured (archipelago-like) and unstructured (fully connected and spatially homogeneous) landscapes differ in richness and spatial species configuration, and what are the relative roles of dispersal heterogeneity and spatial genetic structure in generating these patterns?
(2) Do dispersal evolution, dispersal heterogeneity, and spatial genetic structure alter speciation and extinction dynamics, and thereby the mechanisms maintaining stable species richness?
(3) How does dispersal evolve following speciation, and how does isolation shape this trajectory?

## Methods

### Model overview

We develop a spatially explicit individual-based model (IBM) to examine how dispersal evolution, dispersal heterogeneity, and the spatial genetic structure generated by dispersal inheritance shape speciation and metacommunity dynamics across landscapes. The model builds on established frameworks for understanding sympatric speciation under neutral (non adaptive) evolution (Manzo & Peliti 1994; de Aguiar et al. 2009; Princepe et al. 2022). It operates in discrete, non-overlapping generations and considers spatially distributed individuals on a two-dimensional lattice (Fig. 1).

**Figure 1.**
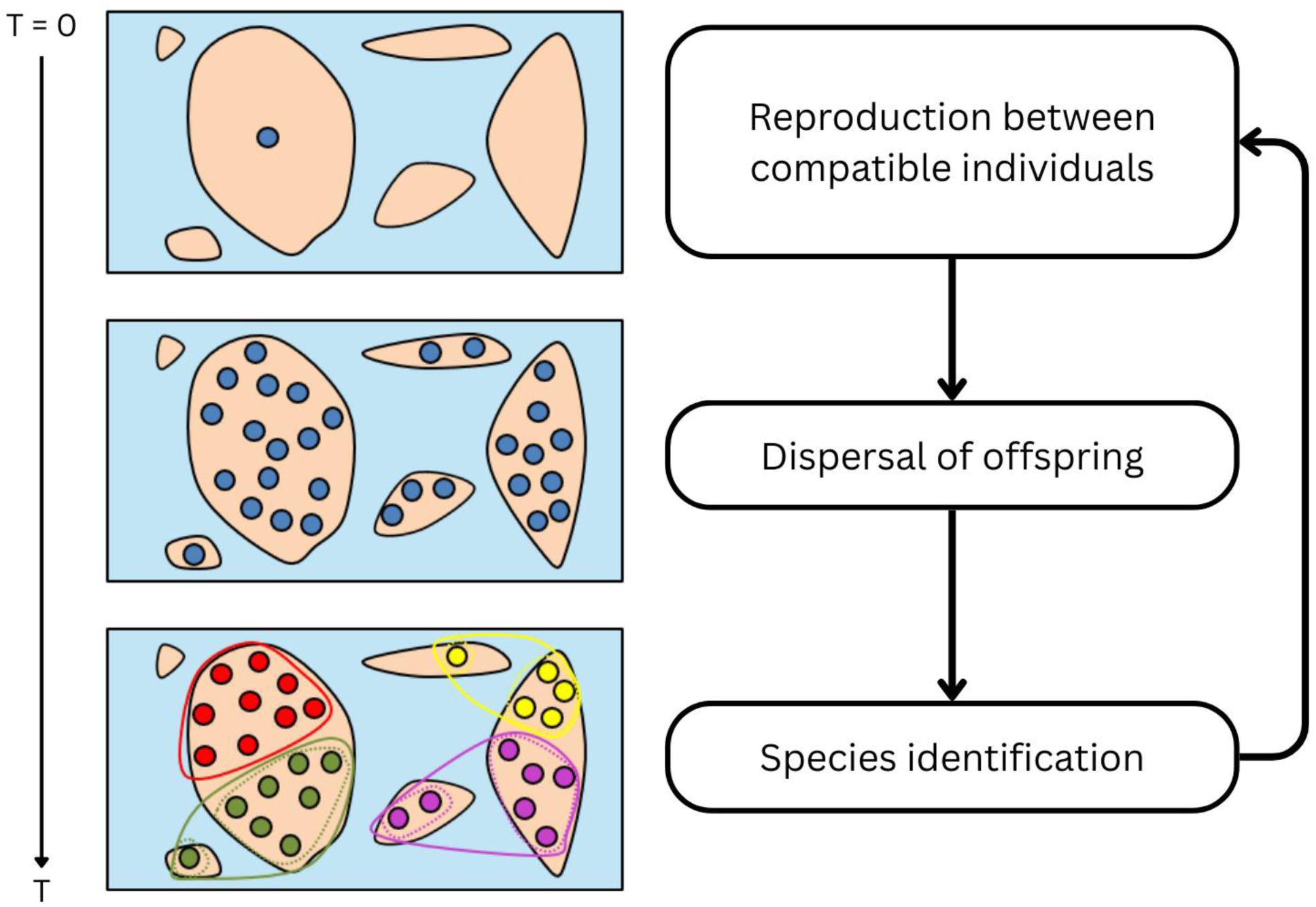
Overview of the individual-based model. Individuals (filled circles) disperse, mate, and reproduce on a two-dimensional lattice (filled rectangle). Colors represent distinct species, identified as genetically cohesive clusters. Polygons illustrate species boundaries that arise from transitive genetic similarity. Each generation includes reproduction, dispersal, local density regulation, and species identification.

### Landscape and population structure

We implement two contrasting spatial configurations to assess how landscape structure interacts with dispersal evolution, dispersal heterogeneity, and spatial genetic structure to shape speciation and metacommunity dynamics. In the structured landscape, the model operates on a two-dimensional square lattice of size *L* = 100, which defines the spatial limits. We first initialize a random field by assigning each grid cell a random value drawn from a uniform distribution between 0 and 1. A Gaussian filter is then applied to the field (*σ* = 4), smoothing local variations by averaging neighboring cell values. This smoothing introduces spatial autocorrelation, such that nearby cells tend to have more similar values, resulting in spatially clustered regions of high and low values. The smoothed field is subsequently converted into a binary habitat map by applying a threshold determined by the desired number of suitable habitat cells (*n*_ℎ*abitat*_ = 0.25 ∗ *L*^2^ for 25% land coverage). Cells with values greater than or equal to this threshold are classified as suitable habitat (land), while cells below the threshold are classified as unsuitable habitat (water). This procedure controls the total amount of suitable habitat while preserving the spatial structure induced by the Gaussian filter. The filter’s standard deviation (*σ*) determines the spatial scale of smoothing and thus influences the size and configuration of habitat patches (islands).

Each island is assigned a “Connectivity Value”. The connectivity value of an island *C_i_* is calculated as the sum of the connectivity *G_ij_* between each pairwise combination of islands, depending on the size of the island itself (*A_i_*) and the island with the nearest point to the focal point (*A_j_*), with *d* representing the smallest distance between these two points.

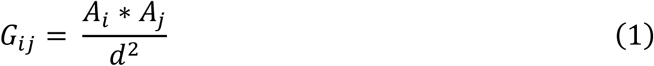

And

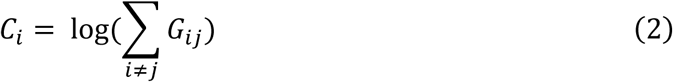

In contrast, the unstructured landscape considers all grid cells a suitable habitat, representing thus a fully connected homogeneous environment with no spatial barriers. To correct for total habitable area, the dimensions of this landscape have been reduced to L/2.

Each scenario is tested under 10 landscape configurations with varying degrees of connectivity and spatial layout, and these are identical among replicates to ensure that replicate runs experience identical underlying spatial structure. At the start of each simulation, a population of 1,000 individuals is initialized in a single grid cell. The initial cell is selected as the location with the highest value in the Gaussian environmental field, representing the most favorable location in the landscape according to the environmental variable used to construct the field. As described above, Gaussian smoothing introduces spatial autocorrelation, whereby neighboring cells tend to have similar environmental values. Consequently, the initial population is surrounded by cells with relatively favorable environmental conditions, potentially facilitating population establishment and subsequent dispersal into adjacent areas. Following initialization, local population dynamics are governed by a carrying capacity of *K* = 2 per grid cell, such that individuals subsequently disperse and no cell can sustain more than two individuals. This grid-cell carrying capacity is consistent with previous spatially explicit individual-based models (Mortier et al. 2019; Masier & Bonte 2020). Movement, reproduction, and lineage diversification thus unfold within landscapes that differ only in their spatial configuration (structured vs. unstructured), allowing direct comparison of how spatial heterogeneity shapes metacommunity- and speciation dynamics.

### Individuals and reproduction

Individuals are represented by haploid genomes encoded as linear binary strings *B* ≡ (*S*_1_, *S*_2_, …, *S_B_*), *S_k_* ∈ {0, 1} of size |*B*| = 100. Reproduction occurs once per generation and is structured by both spatial proximity and genetic similarity (de Aguiar et al. 2009; Princepe et al. 2022). Following the seeker-mate mechanism of de Aguiar et al. (2009), each individual acts as a “seeker” that attempts to mate with genetically compatible partners located within a mating radius of *d*_mating_ = 2 cell (see also Kauai et al. 2023). Compatibility is quantified by the genetic similarity *q_ij_* between individuals *i* and *j* that measures the proportion of loci that are homologous as follows:

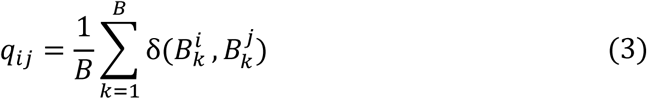

where *δ*(*B_k_^i^,*B*_k_^j^*)=1 if alleles are equal and 0 otherwise. Mating is permitted only when *q_ij_* ≥ *q*_min_ = 0.95, consistent with similarity thresholds used in earlier speciation models (de Aguiar et al. 2009; Princepe et al. 2022, Kauai et al., 2023). Self-fertilization (selfing) is always permitted. In the seeker-mate process, each individual *i* includes itself as a potential mating partner, such that *j* = *i* is always available irrespective of spatial distance or genetic similarity constraints. In this case, genetic compatibility is trivially satisfied (*q_ii_* = 1 ≥ *q*_min_).

Mating proceeds without replacement: once individuals reproduce, they are removed from the mating pool, mimicking monogamous pairing. Offspring genomes are formed by randomly selecting one allele per locus from each parent with equal probability, after which each locus mutates with probability *μ* = 5 × 10^−5^. The number of offspring produced by each successful pair follows a Wright–Fisher process with a mean reproductive output of 3. When selfing occurs, the offspring genome is constructed by copying the alleles from the single parent at each locus, after which mutations are applied with probability *μ* per locus.

At the end of each reproductive iteration, the offspring cohort replaces the parental population entirely, and thus the time-evolution of the system follows discrete non-overlapping generations.

### Dispersal and population dynamics

Following reproduction, each offspring inherits a dispersal capacity that determines its potential movement distance. Offspring dispersal capacity *d*^′^ is drawn from a Gaussian distribution with mean *d*^ˉ^ = (*d_i_* + *d_j_*)/2 and standard deviation *σ* = 3, where *d_i_* is the dispersal capacity of parent *i*. Under simulations with evolving dispersal, this formulation allows dispersal to evolve gradually while simultaneously generating standing variation in dispersal capacity among individuals.

The realized dispersal distance *D* of each offspring corresponds to the distance from the parental seeker to the place where it will become the seeker of the next generation. Realized dispersal is drawn from a Pareto distribution, a fat-tailed kernel widely used to emulate natural dispersal patterns and the occurrence of rare long-distance dispersal (LDD) events (García & Borda-de-Água 2017; Treep et al. 2021). Specifically:

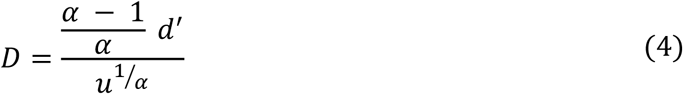

where *u* is a uniformly distributed random number in [0, 1], and *α* = 5, which allows for long distance dispersal (Bullock et al. 2017; Hovestadt et al. 2011). Smaller values of *α* increase the frequency of LDD events, whereas larger values suppress them; sensitivity analyses for *α* are presented in the Supporting Information. The coupling of *d*^′^ with the dispersal kernel ensures that individuals with higher dispersal capacity are more likely to generate long-distance dispersers.

Movement is uninformed and not directed by habitat suitability. After dispersal, density-dependent population regulation occurs independently in each grid cell: if the number of individuals exceeds the local carrying capacity *K*, individuals are removed at random until the limit is reached. We use a constant value *K* = 2 across all suitable cells (both structured and unstructured landscapes), meaning that local density regulation occurs independently within each occupied cell.

To disentangle the relative contributions of dispersal evolution, dispersal heterogeneity, and local spatial genetic structure, we implement three dispersal scenarios. Firstly, we consider a scenario of evolving dispersal, in which dispersal capacity is heritable and subject to mutation as described above. Mutation continuously introduces novel variation, allowing dispersal to evolve while simultaneously generating genetic variance and local spatial genetic structure through inheritance. Secondly, the fixed dispersal scenario comprises of all individuals sharing the same dispersal value throughout the simulation, with offspring inheriting the exact parental trait; this parameterization mirrors earlier speciation models with non-evolving dispersal (de Aguiar et al. 2009) and removes dispersal heterogeneity and prevents the formation of spatial genetic structure associated with dispersal capacity. Finally, we consider a reshuffled dispersal scenario in which, each generation, offspring receive a dispersal value drawn randomly from the population-wide distribution of dispersal, thereby maintaining population-level dispersal heterogeneity while removing local inheritance and the resulting spatial genetic structure associated with dispersal capacity. The three dispersal scenarios constitute a hierarchical decomposition of dispersal processes: fixed dispersal removes heterogeneity, reshuffled dispersal retains heterogeneity but removes inheritance and local spatial structure, and evolving dispersal retains both. Together, these treatments allow us to disentangle the relative contributions of dispersal heterogeneity, dispersal evolution, and spatial genetic structure to colonization, diversification, and long-term biodiversity patterns.

### Species delineation

Identifying species in spatially continuous models poses inherent challenges because boundaries are blurred and divergence proceeds gradually (de Queiroz 2007). To classify individuals into emergent species, we adopt a lineage-based criterion inspired by ring-species systems and compatible with neutral genetic speciation models (de Queiroz 2005; Mallet 1995; Martins et al. 2013). This approach defines species as genetically cohesive units connected through chains of compatible individuals, even when direct mating between all members is not possible.

Genetic compatibility is evaluated using the same similarity threshold applied during mate choice. Two individuals belong to the same species if they are connected by a transitive path of pairwise genetic similarities that are equal to or exceed the minimum threshold *q*_min_ = 0.95. In practice, this means that an individual must share sufficient similarity with at least one other individual in the group, and the entire group must be connected through such links. This criterion ensures that species correspond to cohesive genetic clusters arising from accumulated divergence, consistent with frameworks developed for spatially explicit and individual-based models of speciation (Martins et al. 2013; Rosindell et al. 2010).

To focus on robust speciation events, we exclude ephemeral lineages that persisted for fewer than ten generations. Such short-lived clusters typically reflect transient genetic fluctuations rather than stable divergence, and removing them avoids inflating speciation counts with non-persistent or noise-driven events. For all remaining events, we track lineage identity over time, enabling us to quantify speciation dynamics, extinction events, and the trajectories of dispersal traits associated with each divergence.

### Simulation design and analyses

We conduct simulations across both structured and unstructured landscapes to evaluate the relative contributions of dispersal evolution, dispersal heterogeneity, and local spatial genetic structure to colonisation dynamics, lineage divergence, and long-term metacommunity patterns. For the structured environment, we run ten replicate simulations, thus representing 10 distinct landscape configurations. The unstructured environment is also run for 10 replicates, where each replicate has the identical (homogeneous) landscape. Combined with the three dispersal treatments, this design allows the effects of dispersal heterogeneity and local inheritance to be separated from those of dispersal evolution. All simulations are run for 4000 generations to capture equilibrium-level speciation and extinction dynamics; the first half of each simulation is treated as a burn-in period, and all summary statistics are calculated over the remaining generations.

#### Metacommunity dynamics

For each simulation, we quantify three primary metacommunity metrics: gamma diversity, measured as the total species diversity; beta diversity, measured as the mean similarity in species composition of all pairwise combinations of islands; and degree of island colonization, equaling the percentage of islands habited.

#### Speciation- and extinction dynamics

To relate emergent diversity patterns to the mechanisms underlying dispersal, we examine how speciation and extinction events correlate with island connectivity across the three dispersal scenarios that differ in dispersal heterogeneity and local inheritance.

For each replicate, initial dispersal capacity, generation, and island, speciation events were identified as the first occurrence of each species, whereas extinction events were identified as the final occurrence of each species. Extinction events occurring at the final generation of the simulation were excluded because these may represent species that were still present when the simulation ended rather than true extinction events. The number of speciation and extinction events was then counted for each island and generation.

We modelled speciation and extinction event counts separately using negative-binomial generalized linear mixed-effects models with a log link. For both models, log-transformed island connectivity, dispersal scenario, and initial dispersal capacity, including all interactions, were included as fixed effects (see Supporting Information for model overview). Replicate and island identity were included as random intercepts to account for repeated observations within simulations and differences among islands. Predicted speciation and extinction rates were obtained from the fitted models using marginal predictions across the observed range of log-transformed island connectivity, separately for each dispersal scenario and initial dispersal capacity. Predictions were bias-corrected to account for the nonlinear transformation from the model scale to the response scale and are therefore expressed as the expected number of speciation or extinction events per island per generation.

#### Analysis of dispersal evolution

Opposed to the two previous analyses, dispersal evolution is examined for all speciation events that produced species that persisted for at least ten generations, thereby filtering out ephemeral species. For each event, we extract the mean dispersal capacity of the parent and descendant lineages over time and perform hierarchical clustering using Ward’s method on the resulting temporal profiles. We determine the optimal number of clusters using silhouette analysis (see Supporting Information), which evaluates cluster cohesion and separation across candidate solutions. This analysis allows us to test whether speciation is consistently associated with reduced dispersal, as predicted by island-syndrome theory, or whether multiple post-speciation dispersal trajectories emerge.

## Results

We present the results in three parts. First, we compare metacommunity dynamics between structured and unstructured landscapes under different dispersal scenarios, focusing on species richness and turnover as well as colonization dynamics. Second, we analyze speciation-extinction dynamics between scenarios and across connectivity gradients.

Third, we visualize evolutionary trajectories of dispersal by examining newly formed species and clustering their temporal patterns of trait change.

### General metacommunity dynamics

Landscape structure strongly modifies the effect of the different dispersal scenarios on emergent diversity patterns. In structured landscapes, dispersal heterogeneity, whether generated through dispersal evolution or maintained through reshuffling, substantially increases species richness relative to fixed dispersal scenarios (Fig. 2A). Under fixed dispersal, species richness further increases with increasing dispersal capacity. In contrast, these effects are reversed in unstructured landscapes, where species richness remains comparatively low across all scenarios and reaches its highest values under low fixed dispersal.

**Figure 2.**
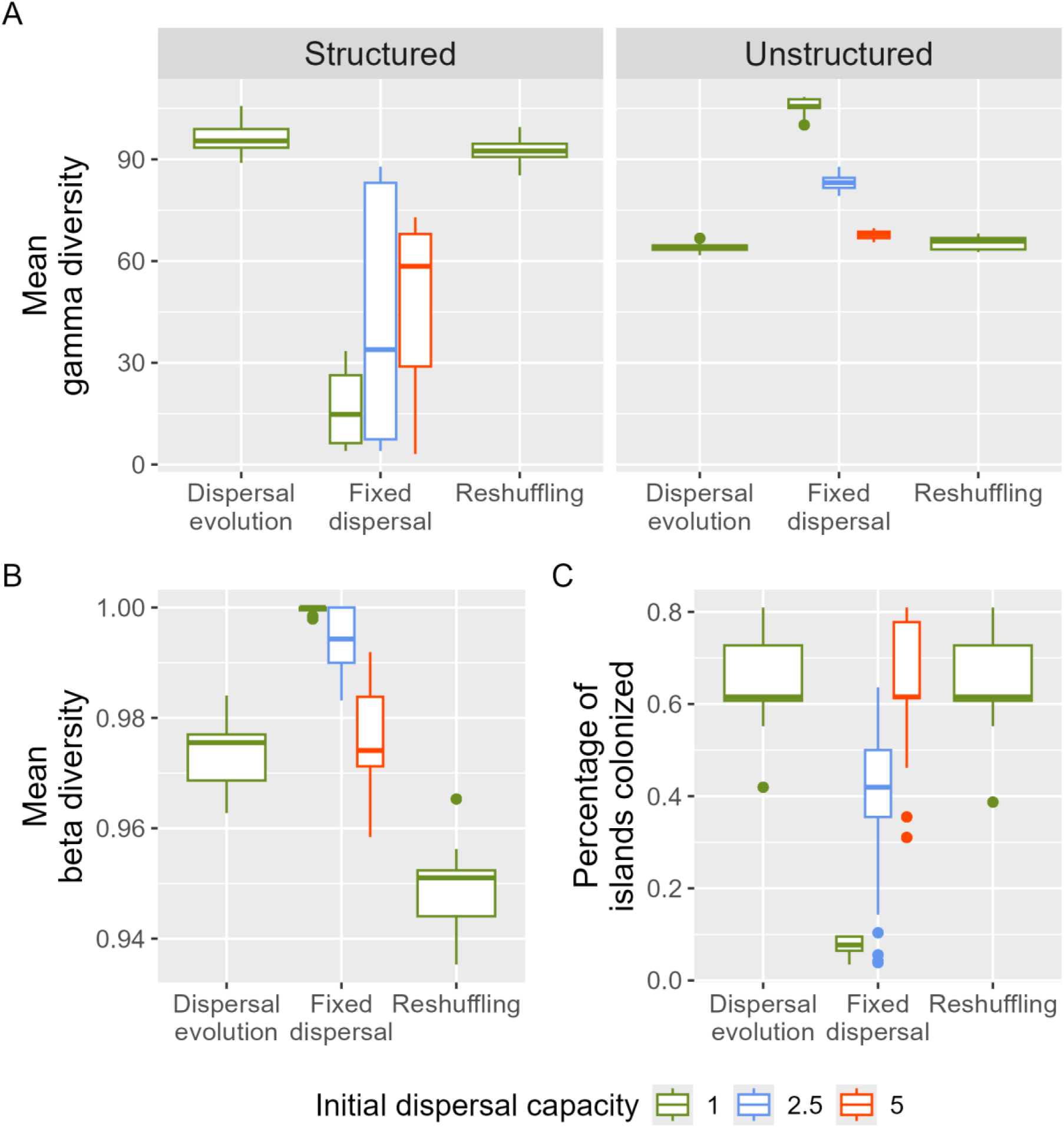
Metacommunity dynamics in unstructured and structured landscapes under different dispersal scenarios. (A) Mean gamma diversity, (B) mean beta diversity, and (C) proportion of islands colonized across dispersal scenarios and initial dispersal capacities. Structured landscapes show greater species richness and spatial turnover under variable dispersal, whereas unstructured landscapes maintain comparatively low diversity. Variable and high dispersal also increase island colonisation, highlighting the interaction between landscape configuration and dispersal in shaping community diversity and spatial differentiation.

Dispersal heterogeneity not only increases species richness in structured landscapes but also promotes greater spatial turnover in species composition (Fig. 2B). Communities with evolving or reshuffled dispersal exhibit stronger differentiation among islands than communities with low fixed dispersal, with the reshuffled scenario showing the strongest effect. At the same time, both heterogeneous dispersal scenarios and high fixed dispersal increase the proportion of islands that become colonized (Fig. 2C), indicating that dispersal heterogeneity expands access to ecological opportunities across the landscape.

The similarity between the evolving and reshuffled dispersal scenarios suggests that many of these biodiversity patterns arise from variation in dispersal capacity itself rather than from dispersal evolution per se. By maintaining a mixture of low- and high-dispersing individuals, heterogeneous dispersal simultaneously promotes colonization of previously unoccupied islands while maintaining opportunities for spatial differentiation among populations.

Together, these results demonstrate that dispersal heterogeneity is a major determinant of metacommunity structure, but that its effects depend strongly on landscape configuration. In unstructured landscapes, high connectivity facilitates widespread gene flow and limits diversification despite increased dispersal. In contrast, structured landscapes allow dispersal heterogeneity to enhance colonization without fully eroding spatial isolation, resulting in greater species richness, stronger community differentiation, and more extensive occupation of available habitat.

### Speciation and extinction dynamics

Despite the strong effects of dispersal scenario on metacommunity structure, speciation and extinction rates remain closely balanced across landscapes and treatments, resulting in relatively stable species richness through time. In structured landscapes, both rates show a weak negative relationship with island connectivity (Fig. 3A), whereas dispersal scenario has a stronger influence on their magnitude. Speciation and extinction rates are highest under low fixed dispersal and lower under both dispersal-heterogeneous scenarios.

**Figure 3.**
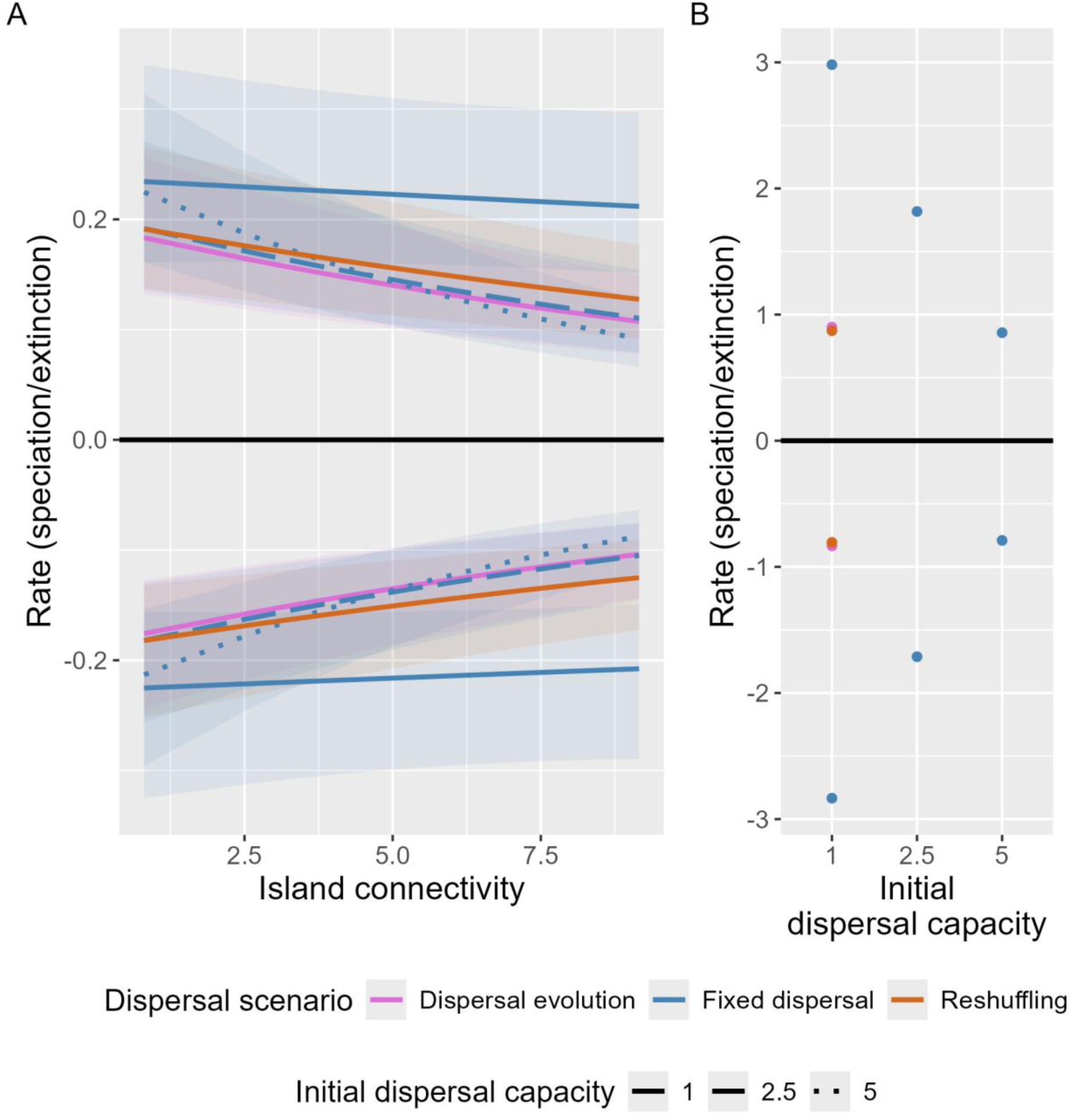
Speciation and extinction as functions of island connectivity. Predicted speciation (positive values) and extinction (negative values) rates across island connectivity for (A) structured and (B) unstructured landscapes. Speciation and extinction rates remain closely balanced across dispersal scenarios, indicating a stable equilibrium in species richness. Rates are generally highest under low fixed dispersal, while increasing island connectivity is associated with a weak decrease in both speciation and extinction rates.

A similar pattern emerges in unstructured landscapes (Fig. 3B), where low fixed dispersal likewise produces the highest speciation and extinction rates. Although absolute diversification rates vary among dispersal scenarios, the close correspondence between speciation and extinction indicates that differences in species richness do not primarily arise from shifts in diversification balance. Instead, the contrasting diversity patterns observed among dispersal scenarios appear largely attributable to differences in colonization dynamics and the spatial organization of populations.

Together, these results suggest that dispersal heterogeneity strongly alters metacommunity structure while having comparatively limited effects on the net balance between speciation and extinction.

### Trends in dispersal evolution

Across structured landscapes, dispersal traits follow two distinct evolutionary trajectories after speciation: dispersal either decreases (cluster 1) or increases (cluster 2; Fig. 4A). In both cases, descendant lineages gradually converge towards dispersal capacities similar to their ancestral values. The relative frequency of these trajectories varies little across the island-connectivity gradient (Fig. 4B). Decreasing dispersal accounts for approximately two-thirds of all speciation events, making it the more common outcome, but increasing dispersal nevertheless represents a substantial fraction of successful divergence events. These results indicate that speciation is not associated with a single characteristic dispersal response. While reduced dispersal occurs most frequently, consistent with expectations that lower connectivity can facilitate divergence, a sizeable proportion of speciation events are associated with increased dispersal. The weak relationship with island connectivity further suggests that local ecological and demographic conditions may be at least as important as spatial isolation in determining post-speciation dispersal trajectories. More broadly, these findings reveal multiple pathways linking dispersal and diversification, rather than a universal tendency towards reduced dispersal following speciation.

**Figure 4.**
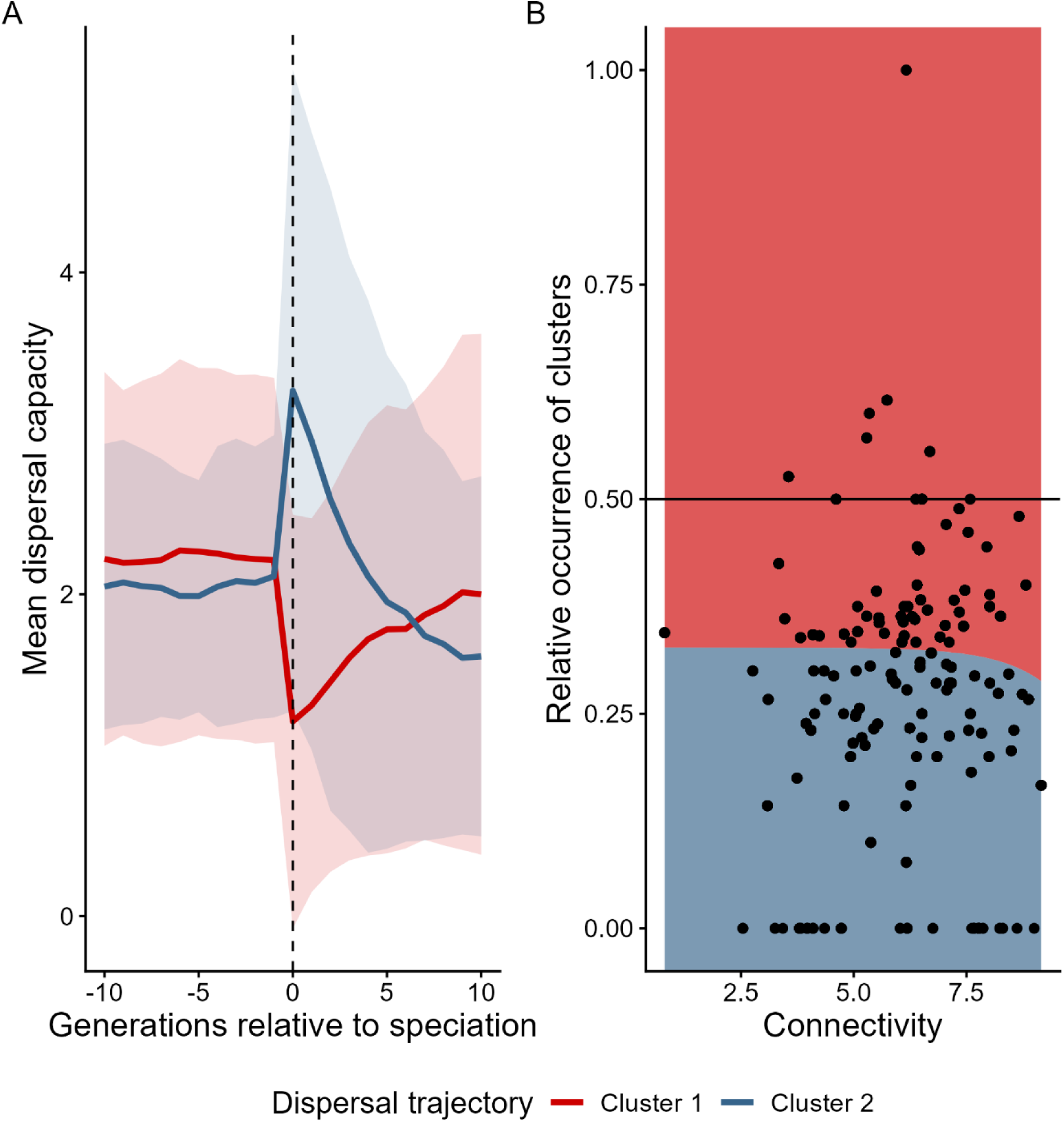
Dispersal evolutionary trends across a connectivity gradient. (A) Dispersal trajectories can be divided into two distinct categories based on their change at speciation: decreasing or increasing dispersal capacity. In both cases, dispersal capacity subsequently converges towards the initial value. (B) The relative frequency of the two trajectories shows no clear association with island connectivity. Decreasing dispersal is the more frequent trajectory, occurring in approximately two-thirds of speciation events across the connectivity gradient.

## Discussion

Our simulations show that dispersal heterogeneity is a key driver of species richness and spatial biodiversity patterns in in spatially structured landscapes. The similarity in metacommunity structure, speciation-extinction dynamics, and colonization patterns between the evolving and reshuffled dispersal treatments, contrasted with their strong differences from fixed dispersal, indicates that dispersal heterogeneity is more important than dispersal evolution itself in shaping long-term biodiversity patterns. Variable dispersal capacity allows lineages to colonize isolated habitats while maintaining local retention, resulting in higher regional diversity than under fixed dispersal. Although isolated habitats are also reached under high fixed dispersal rates, promoting spatial structure, regional diversity does not develop to the same extent. Dispersal variability therefore appears to increase biodiversity not simply by increasing movement, but by generating a broader range of dispersal outcomes: some individuals can colonize distant habitats, whereas others remain locally retained (conform with the findings of Travis et al. 1999). This combination of colonization and retention increases landscape occupancy while maintaining opportunities for spatial differentiation and persistence. Our results therefore suggest that understanding diversification requires considering not only how much organisms disperse, but also how much dispersal varies among individuals. More broadly, these findings reinforce the fundamental role of dispersal in shaping population, community, and diversification dynamics in insular systems (Cowie 2006; Whittaker et al. 2017; Schrader et al. 2021).

A likely explanation for this pattern is that dispersal heterogeneity relaxes the classical trade-off between colonization and retention. Fixed dispersal imposes a single movement strategy on all individuals, forcing a compromise between reaching new habitats and remaining within established populations. In contrast, variable dispersal enables different individuals to occupy different positions along this continuum. Some individuals can reach distant habitats and establish new populations, whereas others experience local retention, maintaining opportunities for spatial differentiation and long-term persistence. This combination of colonization and retention increases landscape occupancy while preserving the conditions necessary for divergence. Understanding biodiversity dynamics in spatially structured landscapes therefore requires considering not only how much organisms disperse, but also how much dispersal varies among individuals. Although the reshuffled treatment represents an artificial manipulation designed to remove inherited evolutionary trajectories, similar heterogeneity is likely common in natural systems. Variation in dispersal behaviour can arise through phenotypic plasticity, condition-dependent dispersal (Clobert et al. 2009), responses to environmental disturbances (O’Connor et al. 2011), interactions with competitors, predators or mutualists (Bestion et al. 2024; Bonte et al. 2024; Fronhofer et al. 2018), and density-dependent dispersal decisions (Bitume et al. 2013). In addition, kin structure and relatedness can strongly influence dispersal strategies, and may become particularly important following founder events or population bottlenecks that reduce genetic diversity (Bitume 2013; Van Petegem et al. 2018). The similarity between the reshuffled and evolving treatments suggests that biodiversity dynamics depend primarily on the presence of dispersal heterogeneity itself, rather than on the particular evolutionary, ecological, or social mechanisms through which that heterogeneity is generated.

These findings extend classical island biogeography theory, which predicts that island size and isolation influence biodiversity through their effects on colonisation and extinction (MacArthur & Wilson 1967). Our results suggest that colonisation success may additionally depend on variability of dispersal capacity. Treating dispersal as a fixed species-level property may therefore overlook the potential for lineages to reach and persist in isolated habitats, as dispersal traits influence both colonization success and subsequent evolutionary dynamics (Alzate et al. 2019; Arendt 2015). This is particularly relevant in structured landscapes, where successful long-distance dispersal can create new populations while limited subsequent dispersal can allow reproductive divergence to accumulate (Gavrilets & Hastings 1996; Feder et al. 2013). Our results therefore suggest that island biogeography may benefit from treating dispersal as a distribution of individual movement capacities rather than as a fixed species-level trait.

Across all scenarios, species richness reaches a dynamic equilibrium in which speciation and extinction balance each other out. Such equilibria are a central feature of biodiversity and metacommunity theory, where long-term richness emerges from the balance between lineage emergence and extinction (Storch & Okie 2019; Reijenga et al. 2021). However, the processes producing this equilibrium differ among dispersal scenarios. Speciation and extinction rates are generally higher under fixed dispersal than under variable dispersal, and connectivity shows a weak negative correlation with both rates. Notably, the higher richness observed under heterogeneous dispersal does not appear to result from a sustained increase in net diversification rates. Instead, it arises from differences in colonization success, spatial organization, and the persistence of lineages across the landscape and thus from how species occupy the landscape rather than from persistent changes in the balance between speciation and extinction.

In particular, dispersal can interact with competition to shape species coexistence and diversity, with evolutionary changes in dispersal potentially altering these relationships (Laroche et al. 2016). Under fixed dispersal, the absence of dispersal heterogeneity may increase competition and turnover by limiting the range of spatial opportunities that individuals can exploit. Dispersal variability may dampen these dynamics by allowing individuals to exploit a wider range of spatial opportunities and persist for longer. Under these conditions, we expect the species richness to be mostly determined by the ecological limit imposed by the environment (Storch & Okie 2019; Wellborn & Langerhans 2015; Yoder et al. 2010). We find that these factors are also influenced by the structure of the landscape: in unstructured landscapes, speciation and extinction occur more frequently compared to structured landscapes. This indicates greater temporal species turnover under these conditions, potentially reflecting the greater difficulty of maintaining persistent populations in a homogeneous environment.

We observed two recurrent post-speciation dispersal trajectories: an initial increase or decrease, followed by a return towards the original value. Contrary to our expectations, the relative occurrence of these trajectories is not strongly influenced by connectivity. We expected dispersal reduction to be more common on isolated islands, consistent with empirical evidence showing reduced dispersal in many island taxa relative to mainland populations (Cody & Overton 1996; Fresnillo & Ehlers 2008; Jessop et al. 2018; Schenk 2013). However, our results indicate that reduced dispersal can also promote divergence in larger and better-connected landscapes. Dispersal reduction has repeatedly been associated with population divergence and speciation across diverse taxa, and can arise in response to both spatial isolation and ecological conditions (Waters et al. 2020). A decrease in dispersal reduces gene flow and can therefore facilitate reproductive isolation, demonstrating that this mechanism is not restricted to highly isolated systems.

The increasing dispersal trajectory represents a complementary pathway to diversification. Increased dispersal allows lineages to reach new and potentially unsaturated areas, where colonisation may establish spatially separated populations that subsequently diverge. Long-distance dispersal is an important mechanism of island colonisation and can generate founder effects, particularly when colonisation involves small numbers of individuals and subsequent isolation (Nathan et al. 2003; Gillespie et al. 2012; Alsos et al. 2015). This resembles founder-event models of speciation, in which long-distance colonisation and isolation contribute to lineage divergence (Gavrilets & Hastings 1996). Although the importance of founder-effect speciation remains debated (Moya et al. 1995; Yeung et al. 2011), our results suggest that the evolution of dispersal traits can facilitate both reduced-gene-flow and colonisation-driven routes to diversification. Rather than favouring a single post-speciation trajectory, diversification in our model appears compatible with multiple evolutionary pathways that generate opportunities for population isolation and species emergence.

The frequent occurrence of decreasing dispersal trajectories may additionally reflect community saturation. In our simulations, much of the available environment is occupied, resulting in competition for limited space. Under such conditions, the benefits of dispersal may decline once suitable habitat is already occupied, whereas remaining in favorable habitat may increase local persistence (Travis & Dytham 1999). Selection on dispersal is therefore likely to depend on ecological opportunity. Increased dispersal may be advantageous during colonisation phases when suitable habitat remains available, whereas reduced dispersal may become advantageous once populations have become established. This is consistent with the broader concept of ecological opportunity, whereby the availability of unoccupied habitat can alter the selective environment and promote diversification (Wellborn & Langerhans 2015; Yoder et al. 2010).

Overall, our results indicate that dispersal heterogeneity represents a fundamental but often overlooked component of biodiversity dynamics and a major determinant of how diversification unfolds across spatially structured landscapes. Spatial structure creates opportunities for population isolation, but variability in dispersal capacity determines how effectively lineages can colonize and exploit these opportunities. While dispersal evolution can contribute to diversification, our comparison between evolving and reshuffled dispersal indicates that many biodiversity patterns emerge from dispersal heterogeneity itself. Dispersal reduction can promote divergence by limiting gene flow, whereas increased dispersal can facilitate divergence by establishing new populations in previously unoccupied areas. Incorporating variation in dispersal capacity therefore provides a more complete understanding of how colonisation, extinction, and speciation interact to generate and maintain biodiversity in island systems and other spatially structured landscapes.

