## Supporting Information for "Dispersal heterogeneity, rather than dispersal evolution, drives biodiversity patterns"

Oikos

#### Sensitivity analysis of the Pareto shape parameter α:

Dispersal capacities are generated from Pareto distributions with mean dispersal capacity $x_{mean} = 1$ and shape parameters $\alpha = 3.5/5/6.5$, using $N = 10{,}000$ random samples for each parameter value (Fig. 1). The shape parameter $\alpha$ controls the spread and tail behaviour of the distribution. Lower values of $\alpha$ produce more strongly right-skewed distributions, with greater variability and a heavier upper tail, increasing the relative probability of very high dispersal capacities. As $\alpha$ increases, the distribution becomes less skewed and more concentrated around its mean, and extreme dispersal capacities become less likely.

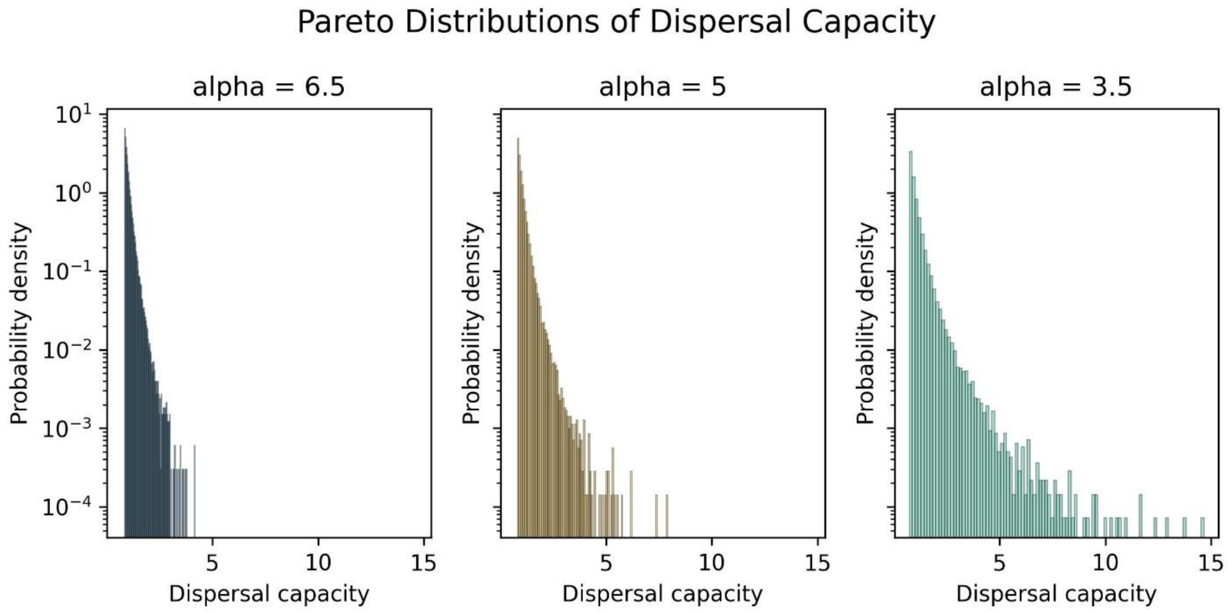


**Figure 1. Distribution of dispersal capacity with multiple α values. Lower values produce more strongly right-skewed distributions and a heavier tail compared to higher values.**

#### Model formulation speciation/extinction dynamics:

Speciation and extinction dynamics are analysed separately for structured and unstructured landscapes using generalized linear mixed-effects models implemented in the R package *glmmTMB*. In both landscape types, the analysis focuses on the second half of the simulation period to characterize dynamics after the initial burn-in phase.

Speciation events are identified as the first recorded occurrence of each species within a simulation replicate and dispersal scenario, whereas extinction events are identified as the last recorded occurrence of each species. Species still present at the final simulation generation are excluded from the extinction counts to avoid classifying extant species as extinct. Event counts are subsequently aggregated by simulation replicate, dispersal scenario, initial dispersal capacity, and generation. For structured landscapes, counts are additionally aggregated by patch, allowing connectivity-dependent variation in speciation and extinction rates to be modelled.

Both speciation and extinction event counts are modelled using negative binomial distributions with a quadratic mean-variance relationship (NB2), which accommodates overdispersion in count data. For structured landscapes, the model includes patch connectivity, dispersal scenario, and initial dispersal capacity as fixed effects, including all two-way and three-way interactions. Random intercepts for simulation replicate and patch identity are included to account for non-independence among observations from the same replicate and patch. The model formulation is:

$$Y_n \sim NB2(\mu_n, \theta),$$

with:

$$\log(\mu_n) = \beta_0 + \log(C_n) * S_n * V_n + b_{replicate(n)} + u_{patch(n)}$$

for the structured landscape, and:

$$\log(\mu_n) = \beta_0 + S_n * V_n + b_{replicate(n)}$$

for the unstructured landscape, with:

- $Y_n$: number of speciation or extinction events
- $\mu_n$: expected event count
- $\log(C_n)$: natural logarithm of island connectivity
- $S_n$: dispersal scenario
- $V_n$: initial dispersal capacity
- $b_n \sim N(0, \sigma^2_{replicate})$: random intercept for simulation replicate
- $u_n \sim N(0, \sigma^2_{patch})$: random intercept for island patch
- $\theta$: negative binomial dispersion parameter, with $Var(Y) = \mu + {}^{\mu^2}/_{\theta}$

Model adequacy is assessed using simulation-based residual diagnostics implemented in the R package *DHARMa*. Model predictions and associated confidence intervals are obtained using the *ggeffects* package, with bias correction enabled. Predicted extinction events are multiplied by (-1) for visualization so that speciation rates appear above zero and extinction rates below zero in the combined figures. This sign change is applied only for visualization and does not affect model fitting.

### Silhouette analysis:

To identify groups of similar trajectories, hierarchical clustering is performed using Euclidean distances calculated between dispersal trajectories. The resulting distance matrix is used to construct a hierarchical clustering dendrogram with Ward's minimum-variance method (ward.D2).

To determine the appropriate number of clusters, candidate solutions with (k=2) to (k=10) clusters are evaluated. For each candidate value of (k), the dendrogram is cut to obtain cluster assignments, and the average silhouette width is calculated using the *cluster* package in R. The average silhouette width measures how well trajectories fit within their assigned clusters relative to neighbouring clusters, with higher values indicating better-defined cluster separation.

The average silhouette width is plotted against the number of clusters, and the number of clusters maximizing the average silhouette width is selected. This analysis identifies two clusters (k=2) as the optimal solution according to the average silhouette criterion.
